# A structural property for reduction of biochemical networks

**DOI:** 10.1101/2021.03.17.435785

**Authors:** Anika Küken, Philipp Wendering, Damoun Langary, Zoran Nikoloski

**Affiliations:** Bioinformatics, Institute of Biochemistry and Biology, University of Potsdam, Potsdam, Germany; Systems Biology and Mathematical Modeling, Max Planck Institute of Molecular Plant Physiology, Potsdam, Germany

**Keywords:** Metabolic networks, network reduction, constraint-based modeling

## Abstract

Large-scale biochemical models are of increasing sizes due to the consideration of interacting organisms and tissues. Model reduction approaches that preserve the flux phenotypes can simplify the analysis and predictions of steady-state metabolic phenotypes. However, existing approaches either restrict functionality of reduced models or do not lead to significant decreases in the number of modelled metabolites. Here, we introduce an approach for model reduction based on the structural property of balancing of complexes that preserves the steady-state fluxes supported by the network and can be efficiently determined at genome scale. Using two large-scale mass-action kinetic models of *Escherichia coli*, we show that our approach results in a substantial reduction of 99% of metabolites. Applications to genome-scale metabolic models across kingdoms of life result in up to 55% and 85% reduction in the number of metabolites when arbitrary and mass-action kinetics is assumed, respectively. We also show that predictions of the specific growth rate from the reduced models match those based on the original models. Since steady-state flux phenotypes from the original model are preserved in the reduced, the approach paves the way for analysing other metabolic phenotypes in large-scale biochemical networks.

## Introduction

Advances in phenotyping, quantitative genetics methods, and systems biology have helped characterize function of genes embedded in different types of biochemical networks ^1–3^. Generation of large-scale stoichiometric models, capturing the metabolism of individual cell types and their interactions in an organism or communities ^1^, has propelled the development of computational approaches for prediction and analysis of metabolic phenotypes ^4^. The sheer size of genome-scale metabolic networks indisputably leads to computational and numerical challenges when employing them to predict and analyse metabolic phenotypes, particularly when these networks are endowed with enzyme kinetics ^5–7^. Against this background we ask: Can large-scale biochemical models be reduced in an unbiased, i.e. fully automated fashion, while still guaranteeing that the steady-state flux phenotypes are preserved in the resulting models of smaller size?

There already exist several frameworks and theories to reduce biochemical models ^8,9^. These approaches can be categorized based on different criteria: (i) whether they only employ the structure of the network (i.e. stoichiometry of biochemical reactions) ^9–13^ or also consider the kinetics of reaction fluxes ^14–23^, (ii) whether they approximate or provide one-to-one correspondence of the steady-state properties between the original and reduced network ^10,23– 25^, and (iii) whether the reduction process is fully automated, i.e. unbiased, or semi-automated (i.e. requires user input) ^9^.

Reduction approaches rooted in the constraint-based modelling framework have been successfully employed to arrive at reduced models that approximate *steady-state flux properties* of large-scale metabolic networks. One group comprises approaches based on user-specified input, consisting of pre-selected metabolites, reactions, and functions (e.g. biomass production) that must be present or supported by the reduced model and, thus, result in reduced models biased by the user’s input ^24–26^. Other approaches in this group start with user-defined subsystems that are subsequently connected (and further refined) to obtain a minimal model that can fulfil a pre-specified function ^27^. In contrast, using ideas from reaction coupling ^28^, a recent unbiased approach provides an efficient reduction of stoichiometric models by reaction removal, while ensuring one-to-one correspondence between the elementary flux modes of the original and reduced model ^10^. Since these approaches do not consider kinetics of reaction flux, they remain silent about the effects that the reduction has on the concentrations of the remaining metabolites.

Approaches for reduction of models with mass action kinetics are particularly important to solve or approximate different reaction mechanisms (e.g. Michaelis-Menten, Hill) ^16,29^. They usually invoke the quasi-steady state assumption (QSSA), whereby the modelled components can be grouped into those with small concentrations, referred to as intermediates, and those with large concentrations (i.e. substrates and products), such that the network displays two (or more) timescales. Invoking QSSA implies that the intermediates, that change on the fast time scale, are expressed as functions of the slow components ^16^. Therefore, these intermediates are eliminated from the original network and, then, the reduced network, capturing the slow dynamics, is further analysed. Application of QSSA is not straightforward since it requires data on kinetic parameters that are challenging to obtain at a genome-scale level ^30^. Therefore, such approaches cannot be analytically carried out ^29^, and often involve approximations that lead to inaccuracies of estimations.

One unbiased technique to eliminate a component in mass action networks requires that the component appears only as a sole substrate or product in the reactions in which it participates^23^. Assuming steady state, the concentration of such a component can be readily expressed as a function of the concentrations of the remaining components. This approach guarantees correspondence between the conservation laws of the original and reduced network and the respective numbers of positive steady states ^23^. This idea has been generalized to allow linear removal of some subsets of components that appear only with stoichiometry at most one in any reaction of a given network endowed with mass action kinetics ^14^. Yet, structural conditions ensuring that the steady-state flux spaces of the reduced network are subset of the reduced network by removal of components of any stoichiometry in networks endowed with arbitrary kinetics remain elusive.

Here we identify such a structural property for model reduction and show that it can be efficiently computed and applied in genome-scale metabolic networks from organisms across the kingdoms of life. The condition allows consideration of biochemical constraints and can, therefore, be invoked with or without assumptions on optimized biological function. The key advantage of the structural condition is that it can be applied to large-scale biochemical networks endowed with mass action kinetics, and under some restrictions, to networks endowed with arbitrary kinetics, while guaranteeing that the steady-state flux phenotypes of the original model are preserved in the reduced.

## Results and Discussion

### Balancing of complexes as a structural condition for network reduction

To illustrate the condition, we first introduce some key concepts from stoichiometric modelling of biochemical networks. A biochemical network is composed of irreversible reactions through which biochemical species acting as substrates are transformed into products. The biochemical network on Fig. 1a is composed of ten reactions that transform six species. The network structure is described by nodes that denote complexes, corresponding to the left- and right-hand sides of the considered reactions, and edges representing the reactions. Therefore, the network on Fig. 1a contains eight complexes connected by ten reactions. The incoming and outgoing neighbourhoods of a complex are given by the complexes to which it is directly connected via incoming and outgoing reactions, respectively. For instance, the incoming neighbourhood of complex 2B is composed of complexes D and F, and the outgoing neighbourhood comprises A+E and F. The stoichiometric matrix of the network, *N*, is then given by the product of the matrix, *Y*, describing the species composition of complexes and the incidence matrix, *A*, of the corresponding directed graph (Methods, Supplementary Fig. S1) ^31,32^. In addition, the reactions are weighted by non-negative numbers which correspond to fluxes of a steady-state flux distribution, *v*, which satisfies *Nv = 0*. The steady-state condition implies that the concentrations of species are invariant in time, i.e. the species are *balanced*, whereby the production and consumption rates are the same.

**Figure 1.**
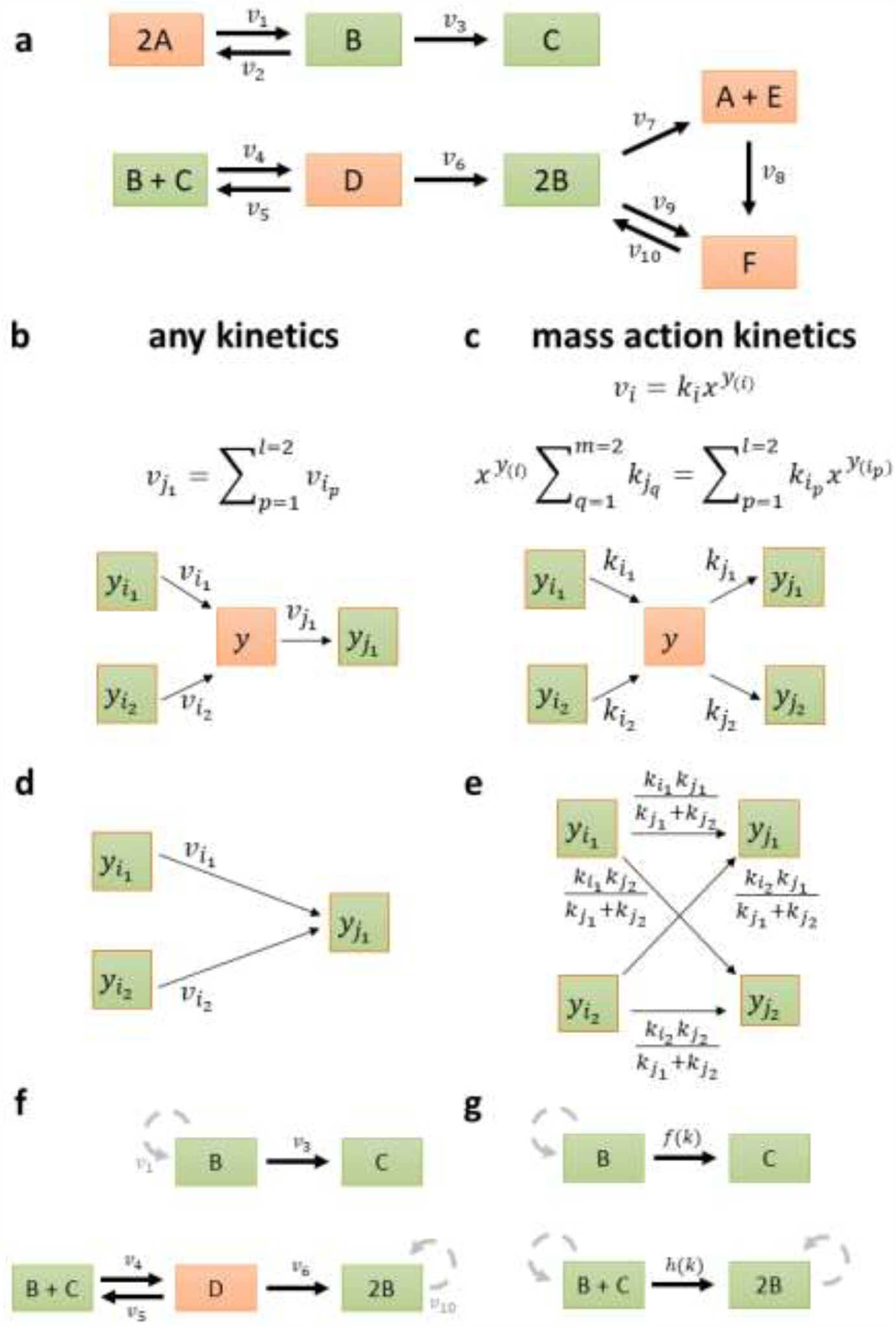
Balanced complexes for network reduction. (**a**) Toy example network borrowed from Shinar and Feinberg ^33^, including six species, A – F, eight complexes, depicted as rectangles, and ten reactions, with rates *v*_*1*_*-v*_*10*_, each connecting two complexes. Balanced complexes are shown in orange, while non-balanced complexes are depicted in green. Balanced complexes 2A, A+E, and F have one outgoing reaction, while the balanced complex D has more than one outgoing reaction. Structural motif allowing removal of the balanced complex *y* (**b**) with single outgoing reaction, for an arbitrary kinetics and (**c**) with multiple outgoing reactions, for mass action kinetics. Removal of the complex amounts to substitution of variables, either with respect to reaction rates *v* or monomials of species concentrations x^y^, which can be represented by (**d**) network rewriting, for arbitrary kinetics and (**e**) additional scaling of reaction rate constants, with mass action kinetics. (**f**) Reduced network obtained by removing the balanced complexes 2A, A+E, and F; the loop reactions are shown in grey since they do not contribute to the stoichiometric matrix. (**g**) Reduced network obtained from the network in panel (**f**) after removal of the balanced complex D, assuming mass action. For simplicity, the rate constants are given as functions *f(k)* and h*(k)*

An analogous condition on balancing can be defined for a single complex: Given a biochemical network, a complex is balanced in a set of steady-state flux distributions *S*. if the sum of fluxes of its incoming and outgoing reactions is the same for every flux distribution in *S*. Clearly, complexes that act as sinks or sources, that only have incoming or outgoing reactions, respectively, cannot be balanced in a network without blocked reactions (i.e. reactions which carry no flux in every steady state the network supports). Further, a complex is considered *trivially balanced* if it includes a species that appears in no other complex in the network. This result is a consequence of the balancing of species due to the steady state assumption. A balanced complex that includes species all of which appear in other complexes are termed *non-trivially balanced*. For instance, complex C cannot be balanced as it is a sink, while D, A+E, and F are trivially balanced since species D, E, and F appear only in these complexes that have both incoming and outgoing reactions. Note that our definition of trivially balanced complexes extends the notion of an intermediate species in other reduction approaches ^23^. Finally, 2A is a non-trivially balanced complex, since A appears in two complexes of which one, A+E, is trivially balanced. In fact, by considering only the steady-state conditions, it is the interlinking of species into complexes that contributes to the formation of balanced complexes.

For a set *S*. of steady-state flux distributions, every balanced complex can be readily identified by constraint-based modelling (Methods). The approach amounts to determining that the minimum and maximum total fluxes around a complex are zero for all steady-state flux distributions in the set *S* These conditions can be verified by linear programming that is efficient even for large-scale networks (Methods). The identification of balanced complexes can be further streamlined by easy screening of trivially balanced as well as sink and source complexes.

Networks in which all complexes are balanced are termed complex balanced, and they can be identified by calculating the structural property of deficiency ^34^. Seminal theoretical results pinpoint that complex balanced networks do not exhibit exotic properties, like multistationarity and oscillations. However, it has not yet been addressed how one can identify individual balanced complexes and what the implications of their presence in the network are with respect to the capacity to exhibit particular phenotypic properties.

We next ask if balanced complexes are relevant for reduction of networks which are not complex balanced. To answer this question, we first consider the scenario with a network endowed with arbitrary kinetics. If a balanced complex has only one outgoing reaction, then the flux of this reaction can be expressed as a sum of the fluxes of incoming reactions. The balanced complex can then be safely removed since the flux of the outgoing reaction can be substituted with the respective sum of fluxes of the incoming reactions for the balanced complex. This is concomitant to introducing a bipartite directed clique, connecting each complex in the incoming neighbourhood with the complex in the outgoing neighbourhood (Fig. 1b,d).. Note that any loop edges introduced by this transformation can be removed, since they do not contribute to the balancing equations. For instance, A+E, F, and 2A can be removed as balanced complexes, resulting in the network on Fig. 1f; however, this cannot be done for the balanced complex D since it has more than one outgoing reaction. Since the removal of complexes in this way results in expressing one flux as a sum of others, the reduced network is unique and does not depend on the order of removing the balanced complexes. A similar idea has been used in the unbiased efficient reduction of stoichiometric models ^10^, however, based on the coupling of reactions around balanced species and without the rewiring of reactions.

Attempting to remove a balanced complex with more than one outgoing reaction fails in general; however, under the constraint that all outgoing reactions are fully coupled (i.e. their flux ratio in every steady state is a unique constant) all outgoing reaction fluxes collapse into one and the balanced complex can be removed. Interestingly, this condition is trivially met for networks endowed with mass action kinetics, since the ratio of fluxes of reactions outgoing from any complex is given by the ratio of the respective rate constants. Importantly, this removal can be carried out even if values for the rate constants are not specified, since the ratio is ensured to be constant. Therefore, under mass action, a balanced complex can be removed by introducing a bipartite directed clique of reactions with rescaled kinetic constants (see Methods, Fig. 1c,e). For instance, the balanced complex D can be removed, like the other balanced complexes in the network, resulting in a mass action network with fewer complexes and species, and reactions with modified kinetic parameters, as shown in Fig. 1g.

The usage of balanced complexes to reduce networks with mass action kinetics generalizes the removal of trivially balanced complexes in which species appear in a single complex with stoichiometry of one ^23^. However, the proposed approach allows removal of species that participate in any number of complexes with any stoichiometry, provided they appear only in balanced complexes. It also provides a sound extension of a reduction approach in which complexes are--arbitrarily--assumed to be balanced ^18^ – and leads to biases in the model reduction since the complexes are not demonstrated to be balanced. In our approach, the conservation laws in the original network hold in the reduced, but not necessarily vice versa (see Methods). Therefore, if the original network has at most one positive steady state for any rate constants and values for the conservation moieties, then the reduced model also has at most one positive steady state for the rescaled rate constants ^23^. Hence, analysis of the reduced model can help identify if multistationarity for the concentration of remaining species is precluded in the original, larger model if seminal, well-established conditions apply ^34^. Note that, the removal of balanced complexes may result in removal of species, but may lead to an increase in the number of reactions. Most importantly, the steady-state concentration of any removed species in the process of model reduction can be expressed as a function of the concentrations of the species in the reduced model (Supplementary Fig. S2).

### Reduction of a large-scale kinetic model of *Escherichia coli* metabolism

To verify that the proposed approach could lead to reduction of metabolic networks endowed with mass action kinetics, we used the genome-scale ^7^ and the so-called core ^35^ kinetic metabolic models of *E. coli*. The core model is composed of 828 species, 1474 irreversible reactions, and 1190 complexes, while the genome-scale model consist of 3002 species, 5237 irreversible reactions, and 4371 complexes. Relying solely on the structure of the network, we identified 484 and 1639 balanced complexes in the core and genome-scale kinetic models, respectively, of which 77.7% and 75.4% were trivially balanced. For instance, the complexes ‘Phosphoenolpyruvate + FBP-Phosphoenolpyruvate -complex’, ‘ACALD-acetaldehyde-NAD-CoA-complex’, and ‘3-phospho-glycerate + PGK-3-phospho-glycerate-complex’ are trivially balanced since the respective enzyme-substrate biochemical complexes appear only in these network complexes. In addition, the complex ‘3-phospho-glycerate + ICL’ is non-trivially balanced, since the species 3-phospho-glycerate appears only in two complexes (‘3-phospho-glycerate + ICL’ and ‘3-phospho-glycerate + PGK-3-phospho-glycerate-complex’ x) and the complex ‘3-phospho-glycerate + PGK-3-phospho-glycerate-complex’ is trivially balanced. From the 484 and 1639 balanced complexes 23% and 26% had only one outgoing reaction; for instance, ‘L-aspartate + PRASCS-ATP-complex’, ‘ATP + PRASCS’, and ‘glucose-6-phosphate + FBP’ are balanced complexes with such properties (see Supplementary Figure S3). The removal of such balanced complexes led to a decrease in the number of species by ∼14% in comparison to both investigated kinetic models. Both models consider elementary reaction steps and, therefore, model species can be of three types: metabolite, enzyme or metabolite-enzyme biochemical complex. We observed that ∼95% of the species removed denoted species that represent metabolite-enzyme biochemical complexes. This is in line with the commonly applied assumption in deriving kinetics that are based on mass action (e.g. Michaelis-Menten), and is due to the fact that metabolite-enzyme complexes, as species, participate in single network complexes that are consequently trivially balanced (as illustrated in the examples above). When considering the additional restriction due to mass action kinetics, we could apply the approach in a second round of reduction. This led to the identification of 694 and 2708 additional non-trivial balanced complexes in the core and genome-scale kinetic model, respectively. Removal of these balanced complexes led to ∼99% decrease in the number **of** species in comparison to the original model for the core as well as the genome-scale model. Therefore, the approach does lead to a substantial reduction in the number of species even when only the structure of the network was employed.

The reduced core kinetic model comprises: fumarate, succinate as well as ubiquinone-8 and ubiquinol-8, interconverted by fumarate reductase (FRD2) and succinate dehydrogenase (SUCDi) together with the related substrate-enzyme complexes (Fig.2). In the reduced genome-scale model extracellular pyruvate, acetate, and L-glucose as well as enzymes exchanging those metabolites with the environment extend the set of species around fumarate reductase and succinate dehydrogenase that are found in the reduced core kinetic model (Fig. 2). Therefore, the steady-state concentrations of all other metabolites in these kinetic models can be expressed as functions of the few species that are present in the reduced model.

**Figure 2.**
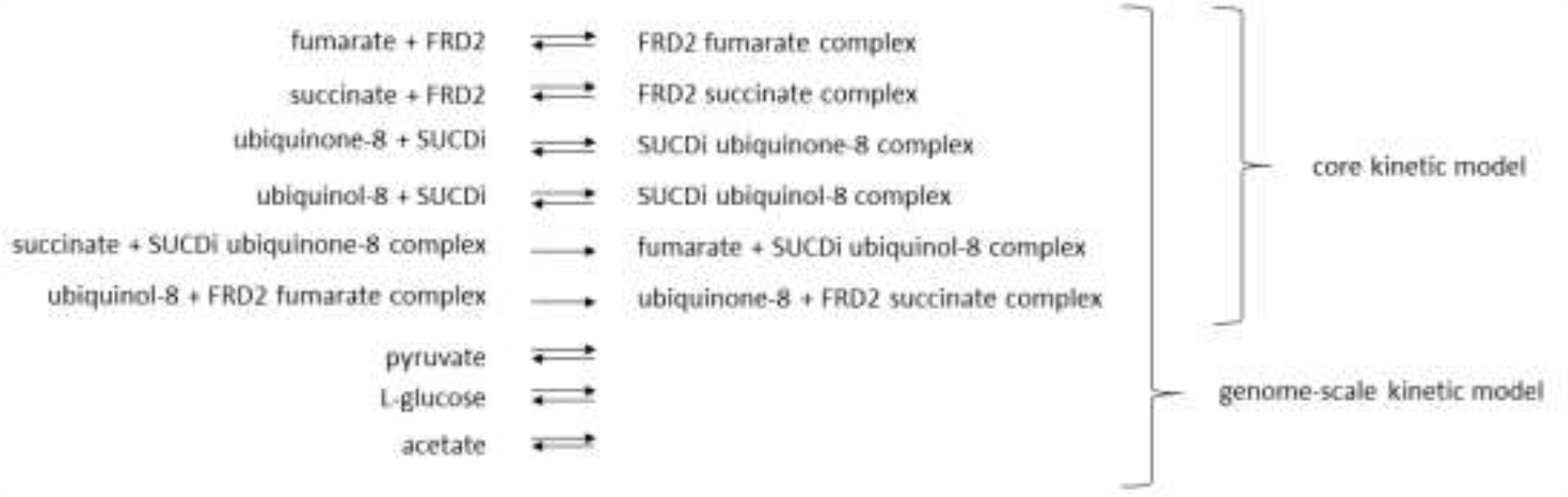
Illustration of the reduced core and genome-scale kinetic models obtained by reduction of a mass action kinetic model. The reduced core kinetic model comprises ten components and ten reactions, while the reduced genome scale kinetic model consists of 13 compounds and 16 reactions. (fumarate reductase - FRD2, succinate dehydrogenase - SUCDi). The concentration of the components present in the reduced networks can be used to model the concentration of all other components in the original networks.

### Reduction of large-scale stoichiometric metabolic networks

Motivated by the findings from the reduction of the core and genome-scale kinetic models of *E. coli* metabolism, we next employed the proposed approach to inspect the reduction in the number of species in genome-scale metabolic models of different organisms. To this end, we used the metabolic networks of twelve organisms to compare and contrast the reductions for the following scenarios: (i) all reactions are assumed to be reversible in contrast to the case when irreversibility constraints, included in the original models, are used, (ii) all reactions follow arbitrary kinetics or their fluxes are described by mass action kinetics, (iii) balanced complexes are determined with respect to the set of steady-state flux distributions compatible with reversibility in comparison to steady-state flux distributions at optimal specific growth rate. Note that reversible reactions are split into two irreversible reactions before applying the approach.

With arbitrary kinetics of reaction fluxes, the general observation was that invoking irreversibility led to only a small increase (< 7%) in the number of removed species across all organisms (Fig. 3a, Supplementary Table S1). The model of *M. barkeri* is an exception to this finding, with 30% increase in species reduction when considering reaction irreversibility. In addition, imposing constraints on biomass had negligible additional effect on the balanced complexes in majority of the networks, except in the models of *P. putida* and *N. pharaonis*, for which there was an increase by 26 and 30% in the number of removed species, respectively (Fig. 3a). This finding demonstrated that the balanced complexes are a property of the network structure and steady-state constraint, rather than due to optimality conditions imposed.

**Figure 3.**
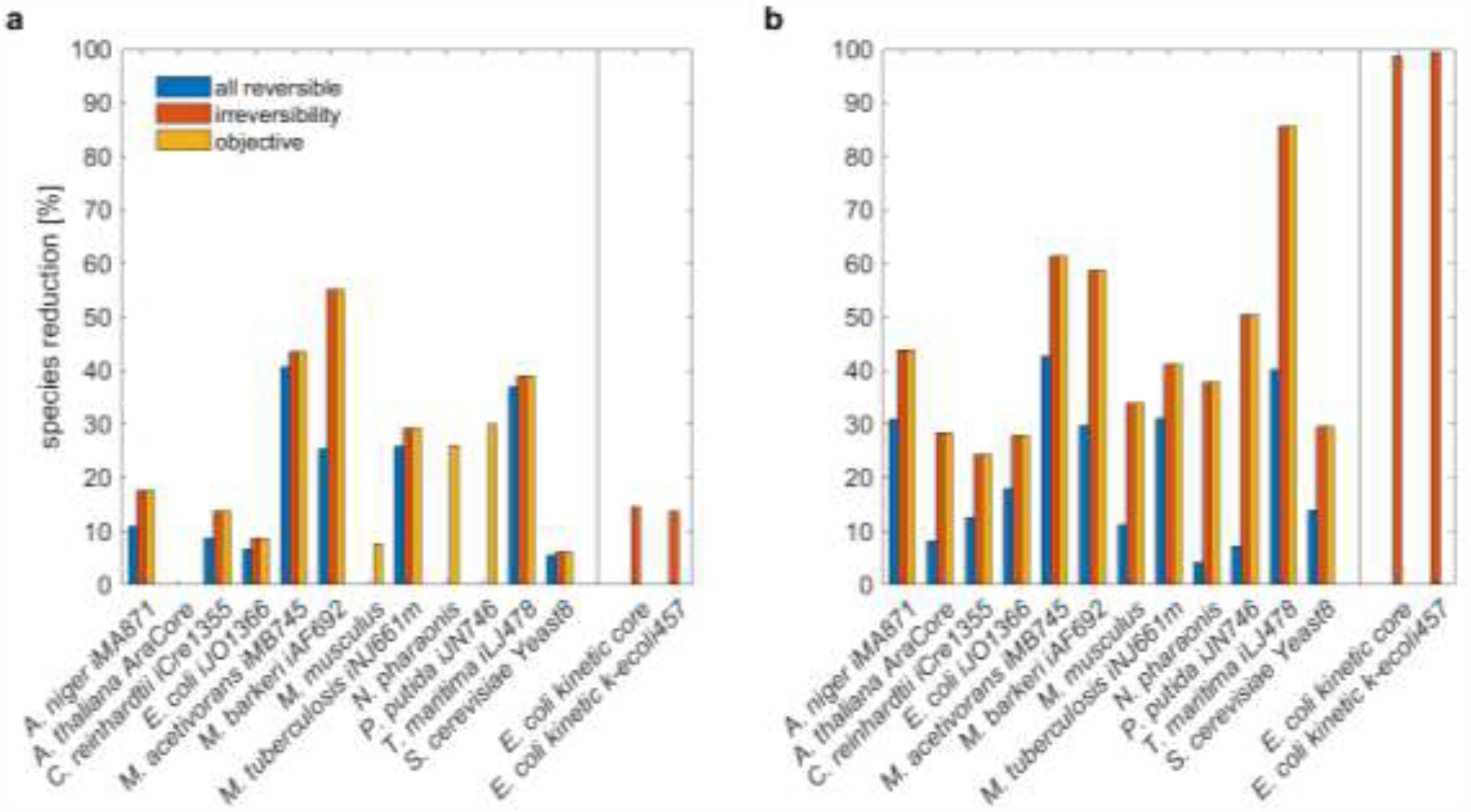
Reduction of genome-scale metabolic networks. Genome-scale metabolic models of twelve organisms from all kingdoms of life are used in the reduction (**a**) with arbitrary kinetics and (**b**) assuming mass action kinetics. The analysis also includes two large, mass-action kinetic models of *E*. coli. Shown is the percentage of reduction in the number of species with respect to the original model.

Consideration of mass action kinetics led to an increase in model reduction of, on average, 26% in comparison to the case of arbitrary kinetics when all reactions are considered reversible (Fig. 3a, b). In contrast to the case of arbitrary kinetics, the assumptions of reaction irreversibility has a large effect on the reduction in the number of species in comparison to the case when all reactions are assumed to be reversible (Fig. 3b). On average, we observed 23% increase in species reduction when reaction irreversibility is considered. Finally, imposing constraints on biomass had no effect on the balanced complexes and, thus, on the number of removed species, when mass action was considered (Fig. 3b).

Altogether, when irreversibility for reactions in the original model was considered, the proposed approach led to no reduction in the number of species in the *Arabidopsis thaliana* core, *M. musculus, N. paharaonis* and *P. putida* model and reached up to 55.1% reduction in the model of *M. barkeri*. On average we observed 18% of species being removed across all considered networks. Instead, when mass action kinetics was considered, we found a reduction between 24.3% for *E. coli* to 85.7% in *T. maritima*, with an average of 44% across the considered models. These results demonstrated that substantial reduction in the number of removed species is possible, while ensuring that the key network properties at steady state are matched between the original and reduced network. In addition, the reduction eliminated up to 17 (12%) species which enter complexes with stoichiometry larger than one (Supplementary Table S2), which is not possible with the existing approaches.

As a general note, the differences in the reduction of the considered models is due to the differences in the number and positioning of balanced complexes with the specific motif of having one outgoing – many incoming reactions (or *vice versa*) in the specific scenarios considered. In comparison to the reduction of the large-scale mass action model of *E. coli*, its counterpart with arbitrary kinetics can be reduced to a smaller degree since its balanced complexes that have the motif structure for reduction are subset of such balanced complexes when mass action is employed. Furthermore, the differences between the scenarios are due to the additional constraints imposed in each of the scenarios.

Next, we investigate compartment-wise species reduction between the original and reduced models, considering reaction irreversibility and mass action kinetics. In doing this analysis, we determined the balanced complexes in the entire network, and then investigated the effect of the reduction by using information about compartments in the respective models. The general observation was that models comprising extracellular space show large reduction for this compartment. The reduction ranged from 26% for *N. pharaonis* to 100% in *T. maritima* (Supplementary Table S3). On average, the reduction over the ten models including extracellular space as a compartment was 65.5%. The average reduction for the cytoplasmic part of bacterial and archaea models was 49%. Compartmented models of *S. cerevisiae, C. reinhardtii* and *A. thaliana* showed largest reduction in mitochondria, chloroplast and cytosol, respectively (Supplementary Table S3).

Using the two genome-scale metabolic models of *E. coli* and *S. cerevisiae* we compared the set of metabolites in the original and reduced models using Jaccard similarity (see Methods). The original models show Jaccard similarity of 0.27 for the set of metabolites, while the reduced models have Jaccard similarity of 0.26. A similar observation was made comparing bacterial models of *E. coli, M. tuberculosis*, and *N. pharaonis* for which at least 95% of the metabolite names could be translated to a common name space. Here, we found Jaccard similarity across the set of metabolites of 0.19 for the original models and 0.16 in the reduced models. These finding demonstrated that the reduction keeps the differences between the original models largely unchanged in comparison to the differences between the reduced models across organisms.

### Predictions of specific growth rate from reduced models

To showcase the benefit of the reduction approach, we considered the reduced models that still include the metabolites (i.e. species) participating in the biomass reactions from the original model (with specified irreversibility). This allows us to compare the simulated specific growth rate between the original and reduced models. The resulting reduced models included 4% fewer metabolites in the model of *A. thaliana* to 28% in the model of *M. musculus* (Fig. 4b). Our findings demonstrate that optimal specific growth rate predicted by the original models are exact match for those of the reduced models (Fig. 4a). Thus, this analysis showcases that the proposed property can be used to remove some well-defined balanced complexes (e.g. those that do not include the building blocks of biomass) without affecting specific growth rates, thus, allowing more targeted applications.

**Figure 4.**
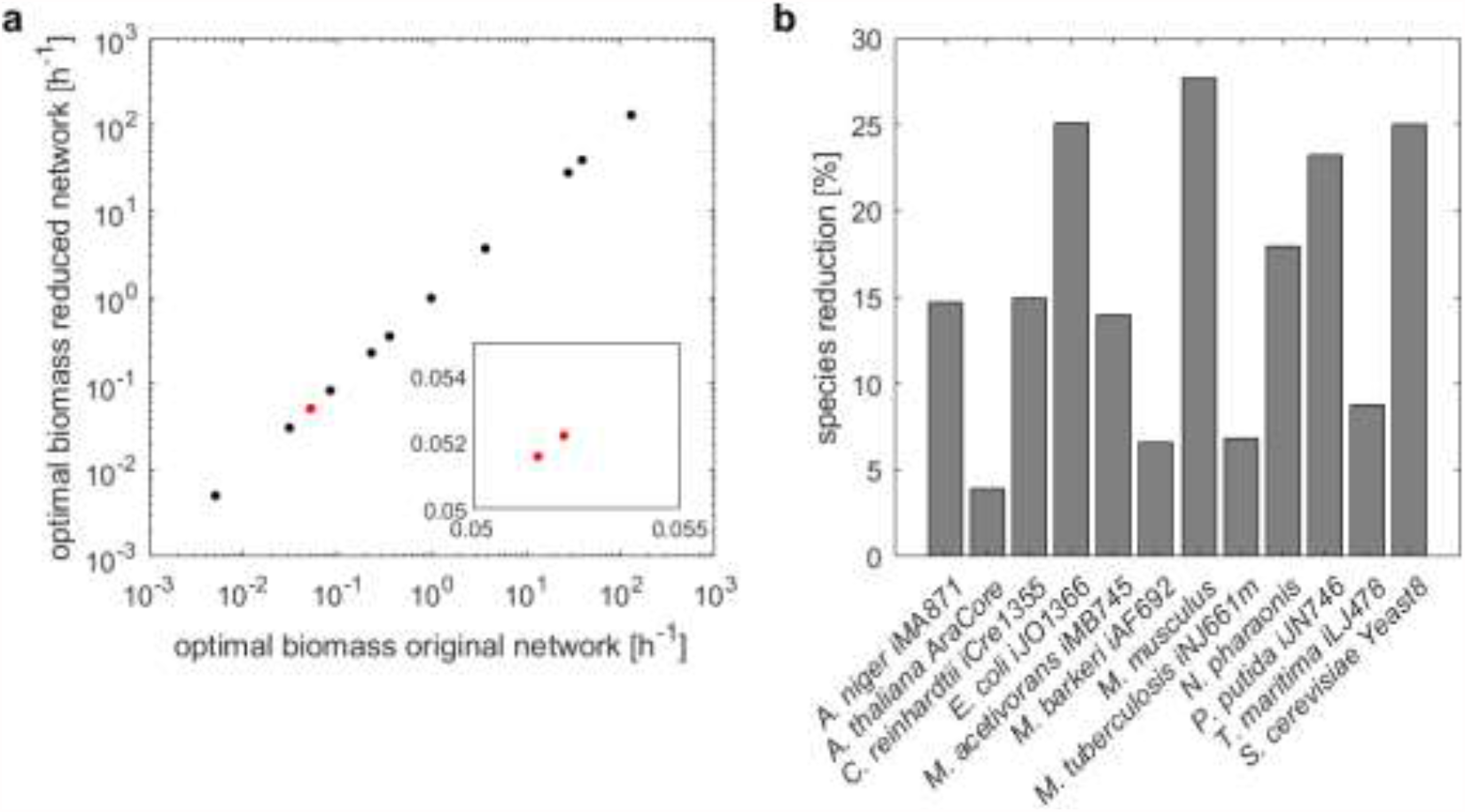
Prediction of specific growth rate in original and reduced models. For direct comparison it is ensured that metabolites that comprise the biomass reaction in the original model remain in the reduced model. (**a**) Specific growth rate estimated from the original and reduced models assuming arbitrary kinetics. The red marked data point are two overlapping points, as shown by the inlay on a smaller scale. (**b**) Model reduction on the level of metabolites.

### Analyses of flux variability and essentiality of reactions

Next, we employed the reduced models used in the prediction of specific growth rates to investigate the difference in the ranges derived from flux variability analysis ^36^. To this end, we calculated the ranges of reactions that are present in the reduced and original model following standard procedure (^36^ Methods). We found that seven of the reduced models showed the same flux ranges at the optimal specific growth rate determined from FBA as those in the original model. In the remaining five models of *C. reinhardtii, M. acetivorans, M. tuberculosis, T. maritima* and *S. cerevisiae*, 71% to 97% of reactions shared between reduced and original model had the same flux ranges at the optimal specific growth rate determined from FBA as those in the original model (Supplementary Figure 4). For the remaining reactions, we observed that in *C. reinhardtii* 94% showed lower flux ranges, while all remaining reactions for *T. maritima* showed larger flux ranges. In models of *M. acetivorans, M. tuberculosis* and *T. maritima* we found that 47% to 76% of the remaining reactions showed larger flux ranges in the reduced models in comparison to the original. The reason why fluxes of some reactions are larger is because the steady state flux distributions of the original network are also steady state flux distributions in the reduced network; however, the reduced network may admit additional steady states (see Methods). These findings demonstrate that the model reduction leaves the flux variability properties (largely) unchanged.

We also inspected the extent to which the reduction affects the predictions of reaction essentiality. To this end, we determined the essential reactions in the reactions shared between the reduced and original models. We found that every reaction that is shared between an original model and its reduced counterpart and is essential in the original model is also essential in the reduced. In addition, the set of reactions introduced during model reduction, e.g. reaction *A* −> *C*,introduced by removal of balanced complex *B* from the path *A* −> *B* −> *C*, contains no essential reactions for the models of *A. niger* and *N. pharaonis*. In contrast, for the models of *M. acetivorans* and *T. maritima* all reactions in these sets are essential. For the remaining models, the percentage of essential reactions in the set of introduced reactions is between 19% and 50%. The essentiality of reactions introduced during model reduction likely results from lumping of at least one essential reaction in the original model. Therefore, we conclude that the proposed reduction does not alter the findings regarding the essentiality of shared reactions and can be used to infer biological role of reactions.

## Conclusion

The constraint-based modelling framework provides a powerful suite of approaches to study the genotype-to-phenotype mapping not only in a single cell, but also across multiple unicellular organisms in a microbial community and interconnected tissues in a multi-cellular organism. However, these advanced applications are associated to increases in the model size which lead to computational and numerical issues in predicting phenotypes of interest. Here, we proposed a fully automated (i.e. unbiased) approach for reduction of models with arbitrary or mass action kinetics at steady state. The approach is based on identifying balanced complexes which contain either a single outgoing reaction, in the case of arbitrary kinetics, or more than one outgoing reaction, for mass action kinetics. We also show that the structural condition can be efficiently identified in large-scale networks, assuming they operate at steady state. Moreover, such balanced complexes can be safely removed from the network as their removal translates into identification of substitution of variables which preserves the steady-state flux distributions of the original model. In addition, the variable substitution can be graphically represented by rewriting the network, leading to a reduced network, and can be carried out without knowledge of kinetic parameters (in the case of mass action kinetics). Most importantly, if a species appears only in balanced complexes, it can be removed from the network and, under the assumption of mass action kinetic, its steady-state concentration can be represented as functions of the steady-state concentrations of the metabolites that remain in the reduced network.

Our extensive analysis of genome-scale kinetic models endowed with mass action show that more than 99% of the metabolites can be removed following the proposed reduction. In addition, since the approach is constraint-based, it also allows us to examine if reaction reversibility or assumption on cellular tasks (e.g. growth) that are optimized have an effect on the size of the reduced model, in terms of number of species. The analysis **of** genome-scale metabolic models across kingdoms of life show that reaction reversibility and assumption of kinetics have the largest effect on the size of the reduced models, leading to up to 85% reduction in the number of species. Nevertheless, we show that if the reduction is performed such that no metabolite participating in the biomass reaction is removed, the reduced model is not affected with respect to predicted growth phenotypes.

We would like to emphasize that once the balanced complexes are identified, it is a choice of the modeller which balanced complexes should be removed in the specific analysis case. For instance, we showed that the removal of the balanced complexes, that do not include species that participate in a biomass reactions, does not affect the prediction of specific growth rates. In this scenario, we also showed that the results from flux variability analysis and predictions of essential reactions in reduced and original models are in very good agreement.

Since the proposed approach is based on a property that can be efficiently determined even in large-scale networks, it provides the means for reduction of multi-tissue and microbial community models. Moreover, it opens the possibility to study the relationship of key properties, such as robustness and multistationarity, between the original and reduced models. However, these applications will have to ensure imposing additional constraints on the removal of the balanced complexes that do not affect the conservation laws in the network. Finally, our approach opens the possibility for unbiased model reduction by seeking to identify other types of structural motifs in large-scale metabolic networks.

## Methods

### Models and their processing

The genome-scale metabolic models of twelve organisms (Supplementary Table S1), were obtained from their original publications ^37–48^. The blocked reactions, reactions with absolute flux values less than 10^−9^ *mmol/gDW/h*, in the metabolic network were determined by Flux Variability Analysis ^49^ and were removed from the original models. Each reversible reaction was split into two irreversible reactions. Two cases for the irreversible reactions originally declared in the model were considered: (1) all were treated as reversible (and were split, as aforementioned) or (2) were maintained as irreversible. The lower bounds for the irreversible reactions were set to zero, while the upper bounds were fixed to the maximum of the upper bounds in the original model. Optimum biomass was determined per Flux Balance Analysis ^50^ with the assumed reaction reversibility.

### Identification of balanced complexes

Let *Y* denote the non-negative matrix of complexes, with rows denoting species and columns representing complexes. The entry *y*_*ij*_ denotes the stoichiometry with which species *i* enters the complex *j* Let *A* denote the incidence matrix of the directed graph with nodes representing complexes and edges denoting reactions. The rows of *A* denote the complexes and its column stand for the reactions. Since the graph is directed, each column of *A* has precisely one -1 and one 1 entry, corresponding to the substrate and product complexes of the respective reaction. The stoichiometric matrix is then given by *N = YA*.

The sum of fluxes around the complex *i* is given by the *i*th entry of the vector *Av*, denoted by [*Av*]_*i*_. A complex is balanced in the set of flux distributions *S ={v* | *NV =0, v*_*min*_ *≤ v ≤ v*_*max*_ *}* if for every *v ∈ S*, it holds that [*Av*]_*i*_ = 0. This condition can be verified by determining that the minimum and maximum values of [*Av*]_*i*_ equal to 0 for each complex *i*, separately. The latter can be ensured by solving two linear programs:

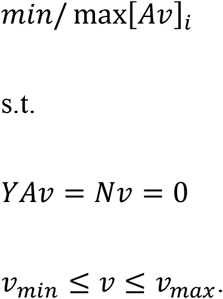

The dependence of balanced complexes on the flux bounds is addressed in follow-up studies^51^.

### Removal of balanced complexes

Suppose that the complex *y* is balanced and that it participates in *l* incoming and *m* outgoing reactions as a product and substrate complex, respectively. Let 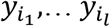and 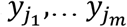denote the substrate and product complexes of the *l* incoming and *m* outgoing reactions.

Let us assume that *m = 1*, i.e., the complex *y* is incident on only one outgoing reaction. Due to the balancing of *y*, 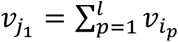. The removal of the complex *y* without affecting the steady-state fluxes entails: (i) removal of the *l + 1* reactions incident on *y* from the original network and (ii) insertion of *l* reactions with a substrate complex from 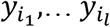 and a product complex 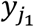. This amounts to substituting every occurrence of 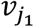 by 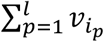, preserving steady-state flux solutions (i.e. the reduced network includes all steady-state flux solutions of the original).

Let us now assume that *m ≥ 1* that the network is endowed with mass action kinetics. The flux of a reaction *i* with complex *y*_*(i)*_ as a substrate is given by 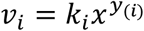, where 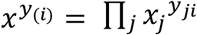 and *y*_*ij*_ denotes the stoichiometric coefficient of species *j* in the complex *y*_*(i)*._ Due to complex balancing, then, 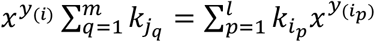. The removal of the complex *y*_*(i)*_ without affecting the steady-state values 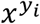 of the complexes entails: (i) removal of the *l + m* reactions incident on *y*_*(i)*_ from the original network and (ii) insertion of *ml* reactions with a substrate complex from 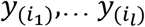and a product complex from 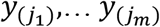. The rate constant of the reaction with 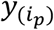 and 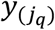 as substrate and product complexes, respectively, is given by 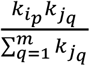. This amounts to substituting every occurrence of 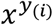 by 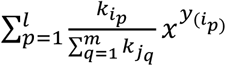 in *Nkφ(x) = 0*, preserving the steady-state flux solutions with respect to*φ(x)* (rather than fluxes, in the case for arbitrary kinetics, above). Note that *φ(x)* is a vector collecting *x*^*y*^ over all complexes, while *k* is the matrix with as many rows as reactions and as many columns as complexes, whose entry *k*_*ij*_ corresponds to the rate constant of the reaction *i* having complex *j* as a substrate or zero, otherwise.

### Preservation of steady states in the reduction process

Let *i* index any species/metabolite and *v* be any steady state flux distribution. It follows from the steady state property *Nv = YAv = 0* that [*YAv]*_*i*_ *= 0;* hence *Σ*_*j*_ *Y*_*ij*_ *[AV]*_*j*_ *= 0*, where *Y*_*ij*_ is the stoichiometric coefficient by which metabolite *i* participates in complex *j*.

Let us consider a single elimination step in this process, in which a particular balanced complex, say *k*, is being removed as explained above. Due to the specific way this removal was defined, the algebraic sum of fluxes for any complex *j* in the reduced network, that is, *[AV]*_*j*._ remains intact, while this sum for the removed complex k, that is, *[AV]*_*k*_ equals zero like that of any other balanced complex. It follows that the steady state equation for metabolite *i*, that is, *Σ*_*j*_ *Y*_*ij*_ *[AV]*_*j*_,= *Σ*_*j≠k*_ *Y*_*ij*_ *[AV]*_*j*_ + *Y*_*ik*_ *[AV]*_*k*_ = 0 still holds after the elimination.

Since the same argument holds for any arbitrary metabolite *i*, it follows that the whole metabolic network will remain at steady state after each removal step. Therefore, any steady state of the original network is conserved at every elimination step and hence, all along the way up to the last reduced model.

### Correspondence of conservation laws

The removal of a balanced complex corresponds structurally to substituting of any directed path of two reactions, *N*_*.i*_ and *N*_*.j*,_ on which the balanced complex is the middle node, with another reaction, given by reaction vector *N*_*.i*_, + *N*_*.j*,_ Clearly, *N*_*.i*,_ + *N*_*.j*,_ *∈ span({N* _*i*,_, *N* _*j*,_*})*. It follows that every new reaction in the reduced network lies in the column span of *N*, Denoting the stoichiometry matrix for the reduced network by <u>*N*</u>, im(<u>*N*</u>) ⊆ im*(N) yields* ker*(N*^*T*^*) ∈* ker*(<u>*N*</u>^*T*^)*. Therefore, for every conservation law, *λ*^*T*^*x = θ* that satisfies *λ ∈* ker*(N*^*T*^), one also has *λ ∈* ker(<u>*N*</u>^*T*^), which means the conservation law is preserved in the reduced network.. However, a conservation law *μ*^*T*^*x = θ* in the reduced network may not necessarily correspond to one of the original network, unless further constraints are imposed on the balanced complexes being removed.

### Predictions of specific growth rates

We calculate optimal specific growth rates by flux balance analysis (FBA) in models with considered irreversibility and compare the predictions before and after removal of balanced complexes. Blocked reactions are removed from the original network and each reversible reaction is split into two irreversible reactions. To avoid cases where no biomass production is ensured in the original model after removal of blocked reactions due to numerical issues, we set the lower bound of the biomass reaction to 5% of the optimal biomass obtained from the model including blocked reactions. In addition, we ensure that the biomass reaction, whose flux is optimized during FBA, is still included in the reduced model. Therefore, we only remove balanced complexes that do not include metabolites (i.e. species) participating in the biomass reactions from the original model.

### Analysis of flux variability and essentiality of reactions

We conducted flux variability analysis ^36^ at 99% of optimal biomass for all reactions that appear in both the original and reduced models considering any kinetic. We then classified the reactions into those that do and do not show differences in the flux ranges between each original and corresponding reduced model. Similar analysis was conducted with respect to reaction essentiality: A reaction that appears in both an original and corresponding reduced model was knocked out and the effect on growth was examined. The reactions were classified as those that are essential or not in each model.

### Data access

The approach is implemented and available together with all data needed to reproduce the findings at https://github.com/ankueken/network_reduction_by_balanced_complexes.

## Acknowledgments

Z.N and D.L are supported by the Max Planck Society.

## Author contributions

Conceptualization: Z.N., data collection: A.K., formal analysis: A.K., investigation: A.K., Z.N., software: A.K., P.W., validation: A.K., writing – original draft: Z.N., writing – reviewing and editing: A.K., P.W., D.L., Z.N.

## References

1. Fang, X., Lloyd, C. J. & Palsson, B. O. Reconstructing organisms in silico: genome-scale models and their emerging applications. Nature Reviews Microbiology (2020). doi:10.1038/s41579-020-00440-4

2. Silverbush, D. & Sharan, R. A systematic approach to orient the human protein– protein interaction network. Nat. Commun. (2019). doi:10.1038/s41467-019-10887-6

3. Thompson, D., Regev, A. & Roy, S. Comparative Analysis of Gene Regulatory Networks: From Network Reconstruction to Evolution. Annu. Rev. Cell Dev. Biol. (2015). doi:10.1146/annurev-cellbio-100913-012908

4. Bordbar, A., Monk, J. M., King, Z. A. & Palsson, B. O. Constraint-based models predict metabolic and associated cellular functions. Nat. Rev. Genet. 15, 107–120 (2014).

5. Stanford, N. J. et al. Systematic construction of kinetic models from genome-scale metabolic networks. PLoS One (2013). doi:10.1371/journal.pone.0079195

6. Chindelevitch, L., Trigg, J., Regev, A. & Berger, B. An exact arithmetic toolbox for a consistent and reproducible structural analysis of metabolic network models. Nat. Commun. (2014). doi:10.1038/ncomms5893

7. Khodayari, A. & Maranas, C. D. A genome-scale Escherichia coli kinetic metabolic model k-ecoli457 satisfying flux data for multiple mutant strains. Nat Commun 7, 13806 (2016).

8. Radulescu, O., Gorban, A. N., Zinovyev, A. & Noel, V. Reduction of dynamical biochemical reactions networks in computational biology. Frontiers in Genetics (2012). doi:10.3389/fgene.2012.00131

9. Singh, D. & Lercher, M. J. Network reduction methods for genome-scale metabolic models. Cellular and Molecular Life Sciences (2020). doi:10.1007/s00018-019-03383-z

10. Tefagh, M. & Boyd, S. P. Metabolic network reductions. bioRxiv 499251 (2019). doi:10.1101/499251

11. Sambamoorthy, G. & Raman, K. MinReact: a systematic approach for identifying minimal metabolic networks. Bioinformatics (2020). doi:10.1093/bioinformatics/btaa497

12. Jonnalagadda, S. & Srinivasan, R. An efficient graph theory based method to identify every minimal reaction set in a metabolic network. BMC Syst. Biol. (2014). doi:10.1186/1752-0509-8-28

13. Clarke, B. L. General method for simplifying chemical networks while preserving overall stoichiometry in reduced mechanisms. J. Chem. Phys. (1992). doi:10.1063/1.463911

14. Sáez, M., Wiuf, C. & Feliu, E. Graphical reduction of reaction networks by linear elimination of species. J. Math. Biol. (2017). doi:10.1007/s00285-016-1028-y

15. Feliu, E. & Wiuf, C. Variable elimination in chemical reaction networks with mass-action kinetics. SIAM J. Appl. Math. (2012). doi:10.1137/110847305

16. Feliu, E., Lax, C., Walcher, S. & Wiuf, C. Quasi-steady state and singular perturbation reduction for reaction networks with non-interacting species. (2019).

17. Radulescu, O., Gorban, A. N., Zinovyev, A. & Lilienbaum, A. Robust simplifications of multiscale biochemical networks. BMC Syst. Biol. (2008). doi:10.1186/1752-0509-2-86

18. Rao, S., der Schaft, A. van, Eunen, K. van, Bakker, B. M. & Jayawardhana, B. A model reduction method for biochemical reaction networks. BMC Syst. Biol. (2014). doi:10.1186/1752-0509-8-52

19. Roussel, M. R. & Fraser, S. J. Invariant manifold methods for metabolic model reduction. Chaos (2001). doi:10.1063/1.1349891

20. Sunnåker, M., Cedersund, G. & Jirstrand, M. A method for zooming of nonlinear models of biochemical systems. BMC Syst. Biol. (2011). doi:10.1186/1752-0509-5-140

21. Danø, S., Madsen, M. F., Schmidt, H. & Cedersund, G. Reduction of a biochemical model with preservation of its basic dynamic properties. FEBS J. (2006). doi:10.1111/j.1742-4658.2006.05485.x

22. Schmidt, H., Madsen, M. F., Danø, S. & Cedersund, G. Complexity reduction of biochemical rate expressions. Bioinformatics (2008). doi:10.1093/bioinformatics/btn035

23. Feliu, E. & Wiuf, C. Simplifying biochemical models with intermediate species. J. R. Soc. Interface (2013). doi:10.1098/rsif.2013.0484

24. Röhl, A. & Bockmayr, A. A mixed-integer linear programming approach to the reduction of genome-scale metabolic networks. BMC Bioinformatics (2017). doi:10.1186/s12859-016-1412-z

25. Erdrich, P., Steuer, R. & Klamt, S. An algorithm for the reduction of genome-scale metabolic network models to meaningful core models. BMC Syst. Biol. (2015). doi:10.1186/s12918-015-0191-x

26. Ataman, M., Hernandez Gardiol, D. F., Fengos, G. & Hatzimanikatis, V. redGEM: Systematic reduction and analysis of genome-scale metabolic reconstructions for development of consistent core metabolic models. PLoS Comput. Biol. (2017). doi:10.1371/journal.pcbi.1005444

27. Baroukh, C., Muñoz-Tamayo, R., Steyer, J. P. & Bernard, O. DRUM: A new framework for metabolic modeling under non-balanced growth. Application to the carbon metabolism of unicellular microalgae. PLoS One (2014). doi:10.1371/journal.pone.0104499

28. Burgard, A. P., Nikolaev, E. V., Schilling, C. H. & Maranas, C. D. Flux coupling analysis of genome-scale metabolic network reconstructions. Genome Res. (2004). doi:10.1101/gr.1926504

29. Pantea, C., Gupta, A., Rawlings, J. B. & Craciun, G. The QSSA in chemical kinetics: As taught and as practiced. in Natural Computing Series (2014). doi:10.1007/978-3-642-40193-0_20

30. Davidi, D. et al. Global characterization of in vivo enzyme catalytic rates and their correspondence to in vitro kcat measurements. Proc Natl Acad Sci U S A 113, 3401– 3406 (2016).

31. Müller, S. & Regensburger, G. Generalized mass action systems: Complex balancing equilibria and sign vectors of the stoichiometric and kinetic-order subspaces. SIAM J. Appl. Math. (2012). doi:10.1137/110847056

32. Neigenfind, J., Grimbs, S. & Nikoloski, Z. On the relation between reactions and complexes of (bio)chemical reaction networks. J. Theor. Biol. (2013). doi:10.1016/j.jtbi.2012.10.016

33. Shinar, G. & Feinberg, M. Structural sources of robustness in biochemical reaction networks. Science (80-.). 327, 1389–1391 (2010).

34. Feinberg, M. Chemical reaction network structure and the stability of complex isothermal reactors - I. The deficiency zero and deficiency one theorems. Chem. Eng. Sci. 42, 2229–2268 (1987).

35. Khodayari, A., Zomorrodi, A. R., Liao, J. C. & Maranas, C. D. A kinetic model of Escherichia coli core metabolism satisfying multiple sets of mutant flux data. Metab. Eng. 25, 50–62 (2014).

36. Gudmundsson, S. & Thiele, I. Computationally efficient flux variability analysis. BMC Bioinformatics (2010). doi:10.1186/1471-2105-11-489

37. Andersen, M. R., Nielsen, M. L. & Nielsen, J. Metabolic model integration of the bibliome, genome, metabolome and reactome of Aspergillus niger. Mol. Syst. Biol. (2008). doi:10.1038/msb.2008.12

38. Arnold, A. & Nikoloski, Z. Bottom-up Metabolic Reconstruction of Arabidopsis and Its Application to Determining the Metabolic Costs of Enzyme Production. Plant Physiol. 165, 1380–1391 (2014).

39. Zhang, Y. et al. Three-dimensional structural view of the central metabolic network of thermotoga maritima. Science (80-.). (2009). doi:10.1126/science.1174671

40. Lu, H. et al. A consensus S. cerevisiae metabolic model Yeast8 and its ecosystem for comprehensively probing cellular metabolism. Nat. Commun. (2019). doi:10.1038/s41467-019-11581-3

41. Imam, S. et al. A refined genome-scale reconstruction of Chlamydomonas metabolism provides a platform for systems-level analyses. Plant J. (2015). doi:10.1111/tpj.13059

42. Orth, J. D. et al. A comprehensive genome-scale reconstruction of Escherichia coli metabolism-2011. Mol. Syst. Biol. (2011). doi:10.1038/msb.2011.65

43. Benedict, M. N., Gonnerman, M. C., Metcalf, W. W. & Price, N. D. Genome-scale metabolic reconstruction and hypothesis testing in the methanogenic archaeon Methanosarcina acetivorans C2A. J. Bacteriol. (2012). doi:10.1128/JB.06040-11

44. Feist, A. M., Scholten, J. C. M., Palsson, B., Brockman, F. J. & Ideker, T. Modeling methanogenesis with a genome-scale metabolic reconstruction of Methanosarcina barkeri. Mol. Syst. Biol. (2006). doi:10.1038/msb4100046

45. Quek, L. E. & Nielsen, L. K. On the reconstruction of the Mus musculus genome-scale metabolic network model. Genome Inform. (2008). doi:10.1142/9781848163324_0008

46. Fang, X., Wallqvist, A. & Reifman, J. Development and analysis of an in vivo-compatible metabolic network of Mycobacterium tuberculosis. BMC Syst. Biol. (2010). doi:10.1186/1752-0509-4-160

47. Gonzalez, O. et al. Characterization of growth and metabolism of the haloalkaliphile Natronomonas pharaonis. PLoS Comput. Biol. (2010). doi:10.1371/journal.pcbi.1000799

48. Nogales, J., Palsson, B. & Thiele, I. A genome-scale metabolic reconstruction of Pseudomonas putida KT2440: iJN746 as a cell factory. BMC Syst. Biol. (2008). doi:10.1186/1752-0509-2-79

49. Gudmundsson, S. & Thiele, I. Computationally efficient flux variability analysis. BMC Bioinformatics (2010). doi:10.1186/1471-2105-11-489

50. Orth, J. D., Thiele, I. & Palsson, B. Ø.O. What is flux balance analysis? Nat Biotechnol 28, 245–248 (2010).

51. Langary, D., Küken, A. & Nikoloski, Z. The unraveling of balanced complexes in metabolic networks. bioRxiv 2021.03.23.436554 (2021). doi:10.1101/2021.03.23.436554

