## Supplementary Figures for "A structural property for reduction of biochemical networks"

**Supplementary Figure**

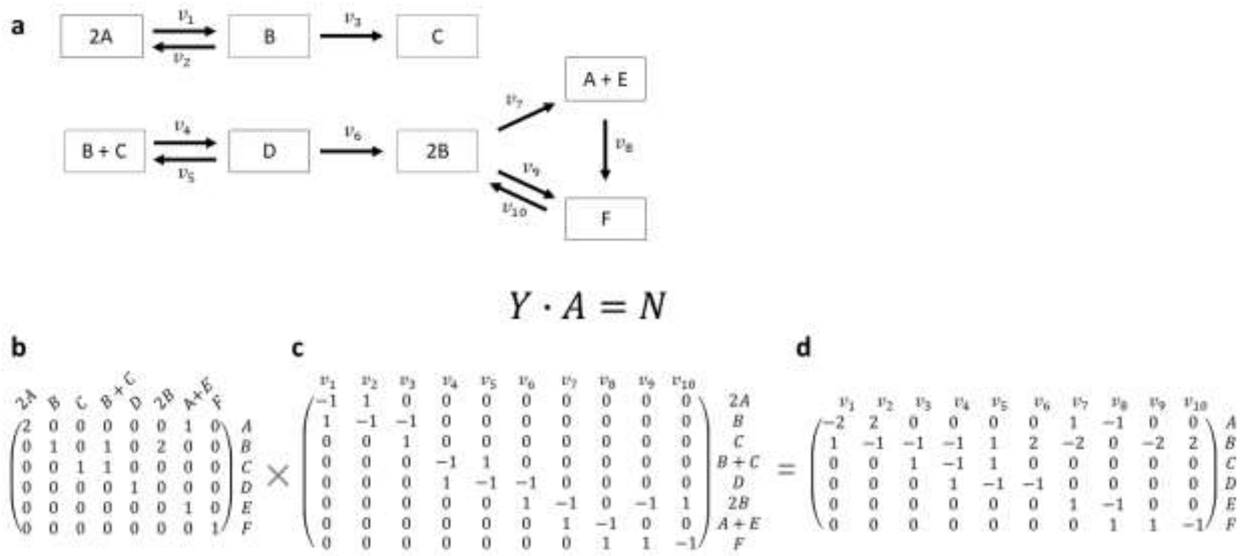


**Supplementary Figure S1. Decomposition of the stoichiometric matrix.** (**a**) Example network borrowed from Shinar and Feinberg (2010), including six species, A – F, eight complexes, depicted as rectangles, and ten reactions, with rates $v_1$-$v_{10}$, each connecting two complexes. (**b**) Species-complex matrix $Y$, where rows correspond to species and column correspond to complexes. Each entry indicates the molarity with which a species participates in a complex. (**c**) Incidence matrix $A$ of the directed graph given in (**a**). (**d**) Stoichiometric matrix $N$, where rows correspond to species, columns correspond to reactions, and each entry indicates the molarity with which a species is produced (positive values) or consumed (negative values) by a reaction. The stoichiometric matrix of the network is given by by the product of species-complex matrix and incidence matrix, $N = Y \cdot A$.

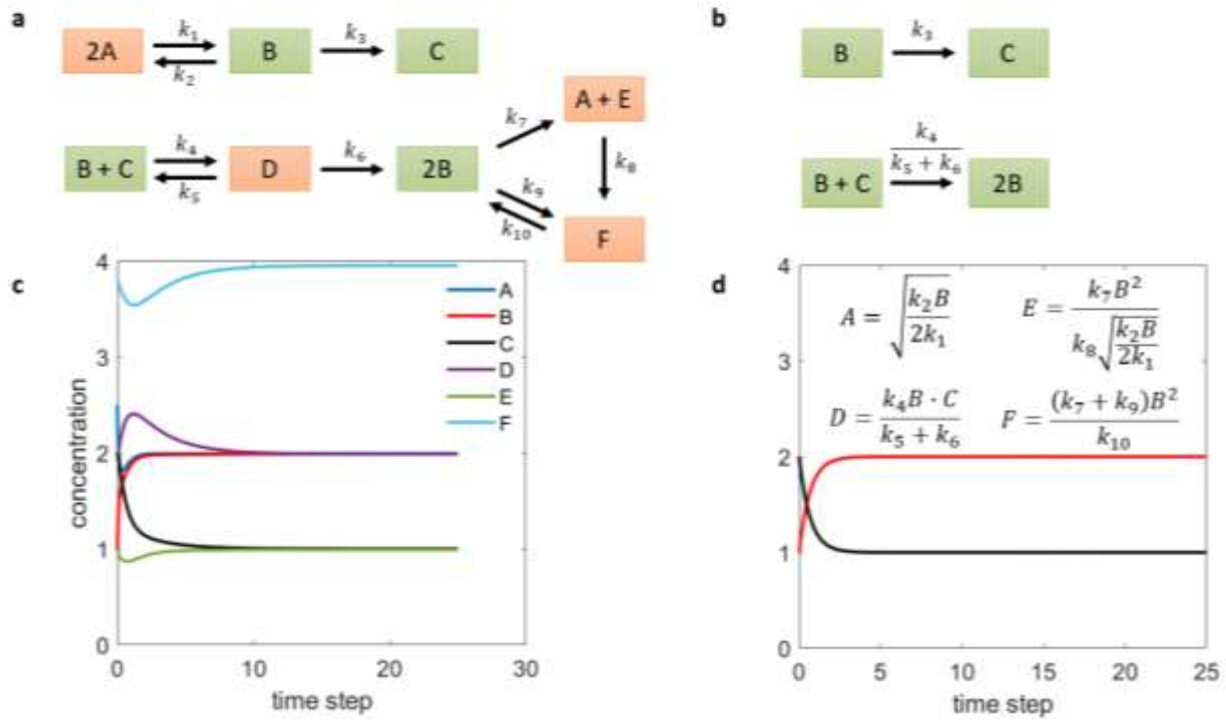


**Supplementary Figure S2. Concentration of removed species can be expressed as function of species in the reduced network.** (**a**) Original network including six species, A – F, eight complexes, depicted as rectangles, and ten reactions, with rates $v_1$-$v_{10}$, each connecting two complexes. (**b**) Reduced network obtained by removing the balanced complexes 2A, A+E, F and D, assuming mass action. (**c**) Simulation of steady-state concentrations for species A – F in the original network using rate constants $k = (1, 2, 0.5, 1, 0.5, 0.5, 0.25, 0.5, 0.25, 0.5)$ and initial concentrations $x = (2.5, 1, 2, 2, 1, 3.8)$. (**d**) Simulation of steady-state concentrations for species B and C in the reduced network using rate constants $k = (0.5, 1)$ and initial concentrations $x = (1, 2)$, like for the original network simulation. Parameters $k$ for the reduced network relate to parameters in the original network as shown in panel (b). Both networks simulate the same steady-state concentrations for species B and C. Concentrations of removed species A, D, E and F can be calculated as functions of species B and C present in the reduced network assuming mass action (see equations in (d)).

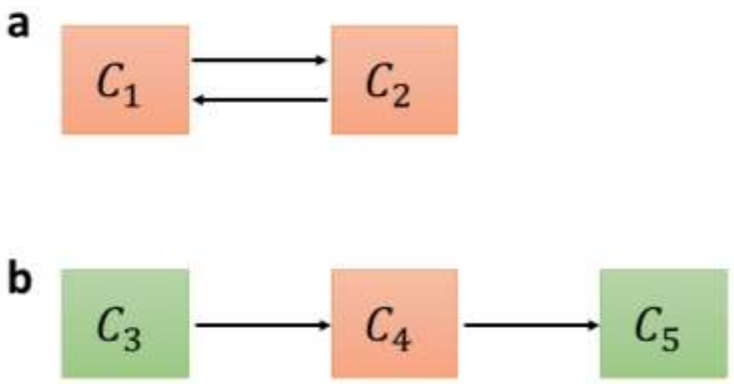


**Supplementary Figure S3. Motifs of balanced complexes with one outgoing reaction in the genome-scale kinetic model of *E. coli*.** (**a**)The motif includes two balanced complexes $C_1$ and $C_2$ which are reversibly converted into each another. (**b**) The motif includes balanced complex $C_4$, its unbalanced substrate complex $C_3$ (having a single outgoing and no incoming reaction) and its unbalanced product complex $C_5$ (having a single incoming and no outgoing reaction).

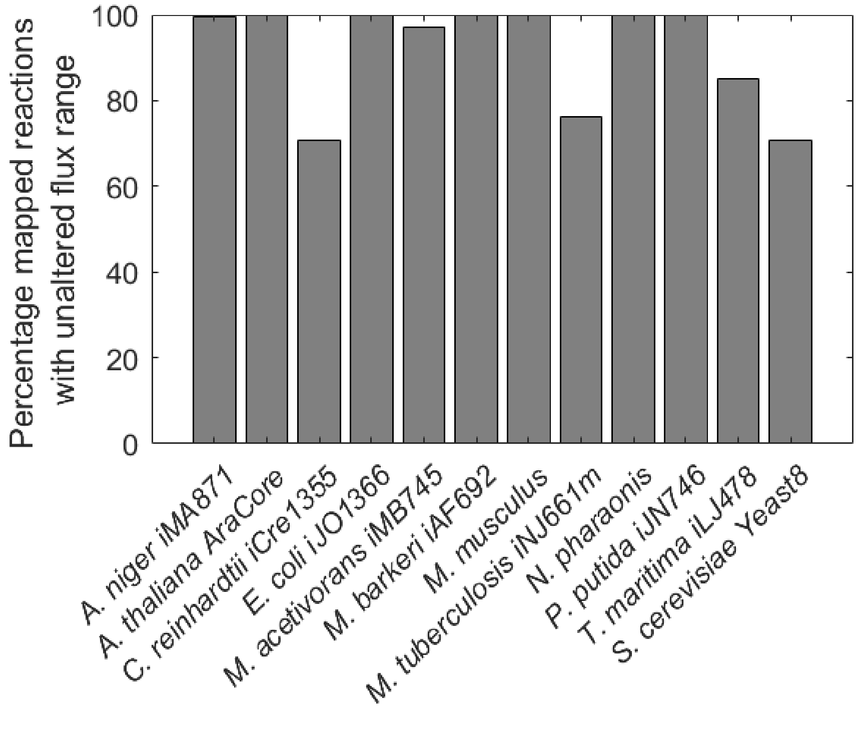

**Supplementary Figure S4. Percentage of reactions unaltered flux range in original and reduced model.** Over the set of reactions with one-to-one mapping between original and reduced model we investigate the percentage of reactions whose flux range obtained from Flux Variability Analysis in the same in the original and reduced model.

### Supplementary Table legends

**Supplementary Table S1. Overview results of balanced complex removal.** The table provides an overview about the number of balanced complexes identified and removed per iteration as well as the type of the balanced complexes (trivial or non-trivial). Moreover, details about the number of species, complexes and reactions in original and reduced models are shown. During analysis three different scenarios were considered (1) every reaction is considered reversible, (2) reaction irreversibility considered, (3) objective (i.e. biomass) optimized. (**a**) Results obtained for the assumption of any kinetics. (**b**) Results obtained for the assumption of mass action kinetics. (**c**) Overview of total model reduction (%) in terms of model species for the different scenarios and assumptions about kinetics (related to Figure 2).

**Supplementary Table S2. Overview of species with stoichiometry larger than one removed from networks during network reduction.** Reaction irreversibility was considered and networks reduced according to the reduction motif of (**a**) any kinetics and (**b**) mass-action kinetics. Metabolite names are provided the way declared in the original publication.

**Supplementary Table S3. Compartment-specific analysis.** Shown are the numbers of metabolites in the original and reduced models for networks of twelve organisms across kingdoms of life.
